# Universality Meets Cultural Nuances: Affective Responses to Rhythmic and Harmonic Complexity Across Cultures

**DOI:** 10.64898/2026.08.31.748217

**Authors:** Mathias Klarlund, Hua Shan, Tomas Matthews, Jan Stupacher, Elvira Brattico, Yi Du, Peter Vuust

**Author notes:** Corresponding author: Yi Du. Equal contributions (Mathias Klarlund, Hua Shan).

## Abstract

The pleasurable urge to move to music—known as groove—has been linked to rhythmic and harmonic complexity via an inverted-U relationship, consistent with predictive coding theory of music. This link has been demonstrated mainly in Western listeners, leaving its cultural generality unclear. Given reports of heightened pitch and harmony sensitivity in Chinese listeners, we predicted stronger responses to harmonic complexity and weaker responses to rhythmic complexity compared with Western listeners. We conducted two online experiments measuring “pleasure” and “wanting to move” ratings for musical excerpts varying in rhythmic and harmonic complexity, comparing Chinese and Danish participants as representatives of Eastern and Western traditions. In Experiment 1, which manipulated both dimensions, both groups showed inverted-U responses to rhythmic complexity, but Danish participants exhibited a sharper curve, indicating greater affective differentiation. No inverted-U pattern emerged for harmonic complexity, and Chinese participants did not show enhanced harmonic sensitivity. Experiment 2 focused on rhythmic complexity using ecologically valid stimuli with finer gradations. Again, Danes showed greater sensitivity to rhythmic variation, reflected in a sharper curvature of their inverted U responses, though peak responses did not differ between groups. Our findings support the universality of predictive responses to rhythmic complexity, while highlighting the modulatory role of cultural exposure in shaping musical affective experiences.

## Introduction

Music is a universal human phenomenon, present in all known cultures (Brown & Jordania, 2013; Nettl, 1999). Yet while musical traditions differ widely, people across cultures often show similar preferences for certain musical features, such as small steps between notes, repeated patterns, and rhythms that fall into regular groupings (Huron, 2001; Savage et al., 2015). Such features are often referred to as *musical universals*, indicating a human tendency to gravitate toward certain types of musical structure. The internalization and processing of such structures and regularities, however, may still depend on cultural exposure and learning (Pezzulo, 2017). While the cultural influences on melody perception has been widely studied (Klarlund et al., 2023; Lahdelma & Eerola, 2020; McDermott et al., 2016; Mehr et al., 2018; Stevens, 2004), less is known about how responses to rhythm and harmony are culturally shaped - despite their central role in musical traditions and their distinct aesthetic principles in Western and Eastern music (Chen, 2018; Nettl, 2005). By comparing Danish and Chinese listeners, as representatives of Western and Eastern musical cultures, this study provides valuable insights into the cultural differences in affective responses to musical complexity.

A commonly observed phenomena in Western music (especially in jazz and funk styles) is groove, namely the pleasurable urge to move to music (PLUMM) or the induction of movements synchronized to an engaging beat (Duman et al., 2024). Empirical aesthetics and music cognition research have established that the groove affective response can be described with an inverted U-shaped relationship between stimulus complexity and positive affective ratings, such as liking, pleasure, and preference(Berlyne, 1971; Witek et al., 2014, 2023) (Berlyne, 1971) (Witek et al., 2023) Witek et al., 2014). More specifically, PLUMM ratings follow an inverted U-shaped relation with rhythmic complexity, typically quantified as syncopation (Matthews et al., 2019; Sioros et al., 2014; Spiech et al., 2022; Stupacher, Wrede, et al., 2022; Witek et al., 2023; Zalta et al., 2024). This relationship is modulated by individual factors like musical training (Matthews et al., 2019, 2022), age (Pando-Naude et al., 2024), Parkinson’s disease (Pando-Naude et al., 2024), and substance use disorders (Stupacher et al., 2025). Moreover, recent work has shown that rhythmic and harmonic complexities interact in shaping PLUMM ratings (Matthews et al., 2019; Stupacher, Wrede, et al., 2022), suggesting that temporal and tonal structures jointly influence PLUMM.

These inverted U effects, as well as inter-individual and inter-population differences, have been primarily interpreted within the predictive processing framework, which posits that affective responses emerge from an optimal balance between predictability and surprise (Stupacher, Matthews, et al., 2022; Vuust et al., 2018, 2022; Witek et al., 2023). In this context, beat and meter are defined as a learned internal model of rhythmic expectations. Rhythmic complexity is often operationalized as the degree of syncopation, which is when a rest or silence on a strong (high probability) metric position is preceded by an onset on a relatively weak (low probability) metric position. Syncopations are therefore considered prediction errors which vary in strength depending on their position in the metric model. Moderately syncopated rhythms allow listeners to infer a metric model that generates relatively strong predictions and thus strongly weighted prediction errors. Conversely, simple rhythms elicit strong predictions but very few predictions errors while highly syncopated rhythms elicit very weak prediction errors due to the uncertainty of the antecedent predictions. This framework suggests that individual and cultural differences in the shape of the inverted U are linked to the relative precision of the metrical model. For example, musicians often exhibit sharper curves which is hypothesized to result from more refined internal models (Matthews et al., 2019; 2020).

Most research on PLUMM has been conducted in WEIRD (Western, Educated, Industrialized, Rich, and Democratic) populations, using Western musical structures that assume internalized metric hierarchies (e.g., strong beats 1 and 3 in 4/4 time). While behavioral evidence supports the influence of metric templates on rhythm perception (Palmer & Krumhansl, 1990), the universality of this process remains untested. Although the predictive processing account of PLUMM assumes universal mechanisms for model learning, the structure of those models can be shaped by informal learning and formal practice (Matthews et al., 2019, 2020, 2023). Thus, it remains unclear whether the inverted U-shaped relation between syncopation (calculated within a Western metric framework) and PLUMM ratings, along with the influence of harmonic complexity, generalizes to non-Western cultures.

In terms of harmony, defined as the simultaneous combination of pitches, cross-cultural studies primarily focused on consonance versus dissonance (Butler & Daston, 1968; Maher, 1976). Western listeners tend to favor certain harmonic intervals and experience harmonic tension differently than listeners from other traditions (Eerola & Lahdelma, 2021; Harrison & Pearce, 2019; McDermott et al., 2016; Milne et al., 2023). An interesting musical culture that differs from the Western oriented ones, is the Chinese. Not only because it concerns circa one fifth of the world population but also because it often draws on the pentatonic scale consisting of five tones versus the diatonic scales of seven tones which are dominant in Western music (Fang et al., 2017; Hang et al., 2022). Scales are the building blocks of all harmony, thus, variation in scale systems may contribute to cross-cultural differences in harmonic perception. Moreover, Chinese individuals have been found to have a higher ability for pitch discrimination as contrasted with Western individuals more skilled with rhythms (Zhang et al., 2020) These findings have been related to language experience: Chinese is a tonal language in which pitch contours convey lexical meaning (Yip, 2002) and this linguistic environment enhances pitch perception in non-verbal domains, including music (Bidelman et al., 2013; Deutsch et al., 2004; Pfordresher & Brown, 2009; Joe et al, 2025, *in submission*). While melody and harmony both rely on pitch, it is unclear whether such pitch advantages extend to harmonic processing. Moreover, Western music often emphasizes metrical regularity and danceable rhythms, whereas traditional Chinese music tends to favor free-flowing, expressive timing that deemphasizes rigid metrical hierarchies (Provine et al, 2002; Levitin et al., 2018; Mok, 1966; Wong, 2012). Taken together, these cultural and cognitive differences motivate a cross-cultural investigation into how rhythmic and harmonic complexity influence affective responses. Here in a first experiment, we examined the interaction between rhythmic and harmonic complexity on pleasure and the urge to move in Chinese and Danish participants. (For consistency, we refer to the latter as *wanting to move* throughout the remainder of the paper). In a second experiment, we focused on rhythmic complexity using a set of Western style drum rhythms with finer-grained variation in syncopation. In both experiments, we expected inverted U-shaped response patterns to musical complexity in Danish and Chinese participants. We also hypothesized that Chinese participants, due to their enhanced pitch perception, would show a different interaction between rhythmic and harmonic complexity than Western listeners. Exploring cross-cultural differences in the well-established inverted U-shaped relationship between musical complexity and affective responses contributes to a more comprehensive understanding of the role of enculturation in predictive processing of music.

## Methods

### Participants

Using G*Power (Faul et al., 2009), according to the 4×4 design in Experiment 1, the minimum total sample size required for ANCOVA between 2 groups with 3 covariates (BMRQ, GMS, MT, as stated below) was calculated as 259 (f = 0.25, α = 0.05, β = 80%). Using this sample size, the power of the linear model in Experiment 2 would be up to 93.1% (23 predictors, α = 0.05, f^2^ = 0.15). Data were collected online from 300 participants, including 149 Chinese and 151 Danish individuals. We excluded 37 Danish participants who were not born in Denmark, 24 participants who failed the attention check (13 Danish participants and 11 Chinese participants), and 5 participants over the age of 40 (3 Danish participants and 2 Chinese participants) to ensure comparable age distributions across groups(Pando-Naude et al., 2024). The final sample comprised 234 participants, consisting of 136 Chinese (55 female, 81 males; M = 22.18 years, SD = 2.41) and 98 Danish individuals (63 female, 35 males; M = 24.93 years, SD = 3.60).

Participants were recruited via the SONA recruitment platform (https://cfin.sona-systems.com), and targeted Facebook groups associated with Aarhus University, Denmark, for Danish participants, and via relevant WeChat groups for Chinese participants. Recruitment advertisements for Danish participants specified that only native Danish individuals fluent in English were eligible; those who did not meet this criterion were excluded from subsequent analyses. The study was approved by the Institutional Review Board at the Danish Neuroscience Centre (DNC-IRB-2023-010).

### Stimuli and Design

#### Experiment 1: Chord-Rhythms

We adopted a chord-rhythm stimulus set encompassing three levels of rhythmic and harmonic complexity (Matthews et al., 2019; Stupacher et al. 2022), and expanded it by introducing a baseline isochronous rhythm and an octave chord (inspired by Pando-Naude et al., 2023). This resulted in a 4 x 4 factorial design (one stimulus per condition, 16 stimuli). Each rhythmic pattern spanned one bar in 4/4 meter, consisted of five onsets per bar (not including the isochronous baseline rhythm, which consisted of four quarter notes), and included an eighth-note hi-hat. The bars were repeated four times at a tempo of 96 beats per minute (bpm), which resulted in a stimulus duration of 10 seconds. Rhythms were derived from *claves,* a fundamental rhythmic pattern in Afro-Cuban and Latin American music (music cultures distinctively different from both Chinese and Western European), with complexity levels derived by the degree of syncopation, calculated according to the method devised by Longuet-Higgins and Lee (1984; see Matthews et al., 2019 and (Fitch & Rosenfeld, 2007), for a detailed description and syncopation values).This approach allows for a systematic manipulation of rhythmic predictability, with medium (original clave), low (clave with shifted syncopated notes to metrically strong beats), and high complexity conditions (increasing syncopation from original by shifting onsets to weaker beats).

The rhythmic onsets consisted of a single chord, which was held constant throughout the 10-second stimulus, to avoid culturally specific harmonic progressions. All chords spanned four octaves within a D major tonality (from D2 to D#5), included six notes, and were presented in piano timbre. The low complexity chord was a simple D major triad in root position and with two inversions. The medium complexity chord added a four-note extension to increase richness while maintaining consonance. The high complexity chord incorporated a fifth interval of a flat ninth, creating significant dissonance. (Harrison & Pearce, 2020; Lahdelma & Eerola, 2020; McDermott et al., 2016)

#### Experiment 2: Drum-Rhythms

Experiment 2 employed 24 drum rhythms composed of bass drum, snare drum, and hi-hat onsets that varied over a large syncopation range. Each rhythm lasted two measures and was repeated twice, resulting in an 8-second duration at 120 bpm. These rhythms were based on real music or didactic drum exercises and were chosen for their relative unfamiliarity and for showing minimal relation between syncopation and the number of onsets. Any original variation in the hi-hat timing was replaced with isochronous eighth-note onsets to standardize temporal resolution across patterns. The syncopation values were calculated using the algorithm from Witek et al. (2014), which adapts Longuet-Higgins and Lee’s (1984) method to polyphonic drum patterns. This algorithm weights the syncopations based on the instrument (bass drum or snare), giving more weight to bass drum than to snare hi-hats (see (Seeberg et al., 2025) for rhythm transcriptions and original sources). We refer to these values as the weighted syncopation index (wSI).

Compared to the chord rhythms, these drum rhythms are more ecologically valid within Western music culture, and cover the syncopation range with higher granularity, thus allowing for analysis of the position of the peak of the inverted U across cultural groups.

### Procedure and Ratings

All participants completed both experiments in a single session, in the same order (experiment 1 followed by experiment 2) via online experiment platform *Gorilla Experiment Builder* (www.gorilla.sc; Anwyl-Irvine et al., 2020).

Participants first provided informed consent and completed a demographic questionnaire including age, gender, nationality, and country of current residency. They then rated their headphone quality and adjusted sound levels to ensure optimal audio presentation. Across the two experiments, participants were presented with 16 chord-rhythm and 24 drum-rhythm stimuli, each presented twice, totaling 80 trials. For each trial, participants provided one of two ratings: *“Pleasure”*, indicating the degree of pleasure the rhythm evoked, and “*Wanting to Move”*, reflecting the extent to which the rhythm made them want to tap their foot, bob their head, or dance. Stimulus order was randomized for each participant. Stimulus sets (chord-rhythms or drum-rhythms) were presented in separate blocks to ensure both ratings were recorded for each stimulus type in sequence. Ratings were provided using a continuous slider ranging from 0 (“none” / “not at all”) to 100 (“very much”), with only descriptive labels visible to participants, without numerical indicators.

To ensure participant engagement and data quality, two attention check trials were included (one per experiment). Each attention check involved a native language speaking voice instructing participants to move the slider all the way to the right (“very much”). Participants were made aware of the attention checks at the start of the experiment to motivate engagement. Between the two experiments (chord-rhythms and drum-rhythms), participants completed the Musical Training and general sophistication subscales of the Goldsmith Musical Sophistication Index (Gold-MSI) (Li et al., 2024; Müllensiefen et al., 2014) and the Barcelona Music Reward Questionnaire (BMRQ) (Mas-Herrero et al., 2013; Wang et al., 2023), both in culturally validated versions for Danish and Chinese participants, respectively. These measures allowed us to control for individual difference in musical training (MT), general musical sophistication (GMS), and musical reward sensitivity.

### Statistical Analysis

#### Exclusion of participants

As online experiments suffer from lower engagement (Rodd, 2024), we applied additional exclusion criteria beyond attention checks. We calculated the standard deviation of each participant’s ratings for each of the four tasks (‘pleasure’ and ‘want to move’ ratings for both chord- and drum-rhythm stimuli). Participants with variability below two standard deviations of the group mean (97.5% one-tailed lower threshold) on any one task were excluded (**Figure S1A & B**). This exclusion criterion was further validated through visual inspection of the excluded participants’ data (**Figure S1C & D**). Consequently, 10 Chinese participants and 1 Danish participant were excluded, resulting in a final sample of 223 participants for subsequent analyses.

#### Control variables

For a descriptive account of the cross-cultural differences on each control variable, we conducted a series of Wilcoxon rank-sum tests. The results indicated that Chinese participants scored significantly higher than Danish participants in GMS (*W* = 7226.50, *p* = 0.020, *r*_pb_ = 0.16) and BMRQ (*W* = 9579.00, *p* < 0.001, *r*_pb_ = 0.49), while no significant difference was found in MT (*W* = 5938.00, *p* = 0.717) (**Figure S2**).

#### Analysis of wanting to move and pleasure ratings

To test our hypotheses about how perceived rhythmic and harmonic complexity interact with culture to shape the affective responses of wanting to move and pleasure, we employed linear mixed-effects modelling using the lme4 package in R (Bates et al., 2015), with post-hoc tests conducted using the emmeans package (Lenth, 2024), with FDR correction applied to ANOVA results and Bonferroni correction used for pairwise contrasts. Model diagnostics (residual normality and homoscedasticity) were assessed using the ggplot2 package (Wickham, 2016).

Given known cross-cultural difference in rating styles (Heine et al., 2002), and our observation that Chinese participants gave generally higher ratings than Danish participants (**Figure S3**), we standardized all ratings (z-scored) within each nationality to retain relative difference between stimuli while controlling for cultural rating biases.

Furthermore, we included MT, GMS, and BMRQ scores as covariates in our models. Because MT and GMS were highly correlated (r = 0.84), we regressed GMS on MT and used the residuals (i.e., variance in GMS not explained by MT) in subsequent models to avoid multicollinearity.

#### Chord-Rhythms

For chord-rhythm stimuli, pleasure and wanting-to-move ratings were entered as dependent variables in separate linear mixed-effects models. Fixed effects included rhythmic complexity, harmonic complexity, and their interactions. Both predictors were polynomial-coded (linear, quadratic, cubic) to test for linear and non-linear effects, as each had four discrete levels.

We also included nationality, MT, residualized GMT and BMRQ scores as covariates, along with their interactions with rhythmic and harmonic complexity. Furthermore, interactions between nationality and each of the other three covariates were included, as were higher-order interactions with rhythmic and harmonic complexity measures, resulting in models with up to four-way interactions. Random effects included by-participant random intercepts and random slopes for rhythmic and harmonic complexity, consistent with our design and modeling strategy.

##### Drum-Rhythms

For drum-rhythms, which varied only in rhythmic complexity (no harmonic manipulation), we also conducted linear mixed-effects models to analyze pleasure and wanting-to-move ratings. Here, rhythmic complexity was operationalized using the wSI. We modeled both linear and quadratic terms of wSI, along with their interactions with nationality, to capture potential inverted-U relationships.

Random effects included by-participant random intercepts and random slopes for both linear and quadratic terms of wSI. Control variables (MT, residualized GMS, and BMRQ) were included in both models.

To evaluate the fit of the quadratic models, we compared them with linear models including only the main and interaction effects of wSI and nationality using likelihood ratio tests and AIC values.

To examine cross-cultural differences in the shape of the inverted-U curves, we conducted a 1,000-iteration bootstrap procedure. In each iteration, we resampled with replacement from the Chinese and Danish participant groups, ensuring sample sizes equal to the original data, and limiting the number of repeated samples to no more than 50 per resample. We then fit a simplified model without covariates to each resampled dataset and extracted the peak (vertex) points of the fitted quadratic curves for each nationality. The difference in peak wSI values between groups was computed for each iteration, and statistical significance was assessed by calculating the 95% confidence interval of the resulting distribution.

## Results

### Experiment 1: Chord-Rhythms

#### Wanting to move

Using wanting-to-move ratings as the dependent variable, linear mixed-effects models (**Table S3**) revealed significant interaction between nationality and rhythmic complexity (*F*_(3, 215.16)_ = 12.13, *q* < 0.001), as well as between nationality and harmonic complexity (*F*_(3, 442.70)_ = 15.03, *q*< 0.001). There were no significant interactions between rhythmic and harmonic complexity (*q* = 0.242), nor a three-way interaction with nationality (*q* = 0.821).

Post-hoc analyses comparing group differences in the contrasts between consecutive levels of rhythmic complexity revealed that Chinese participants had significantly smaller contrasts than Danish participants between the Low and Medium levels (CN: *t*_(215)_ = -1.07, *p*_bonferroni_ = 0.862; DK: *t*_(215)_ = -8.58, *p*_bonferroni_ < 0.001; CN-DK: *t*_(215)_ = 5.33, *p*_bonferroni_ < 0.001), as well as between the Medium and High levels (CN: *t*_(215)_ = 7.97, *p*_bonferroni_ < 0.001; DK: *t*_(215)_ = 15.44, *p*_bonferroni_ < 0.001; CN-DK: *t*_(215)_ = -5.33, *p*_bonferroni_ < 0.001). No group difference was observed for the contrast between the Isochronous and Low levels (CN: *t*_(215)_ = -6.17, *p*_bonferroni_ < 0.001; DK: *t*_(215)_ = -6.40, *p*_bonferroni_ < 0.001; CN-DK: *p*_bonferroni_ = 0.925). These results indicate an inverted-U shape relationship between rhythmic complexity and wanting-to-move ratings, with medium rhythmic complexity eliciting the highest ratings and a more pronounced peak at the medium level for Danish compared to Chinese participants (**Figure 2A**, left panel).

**Figure 1.**
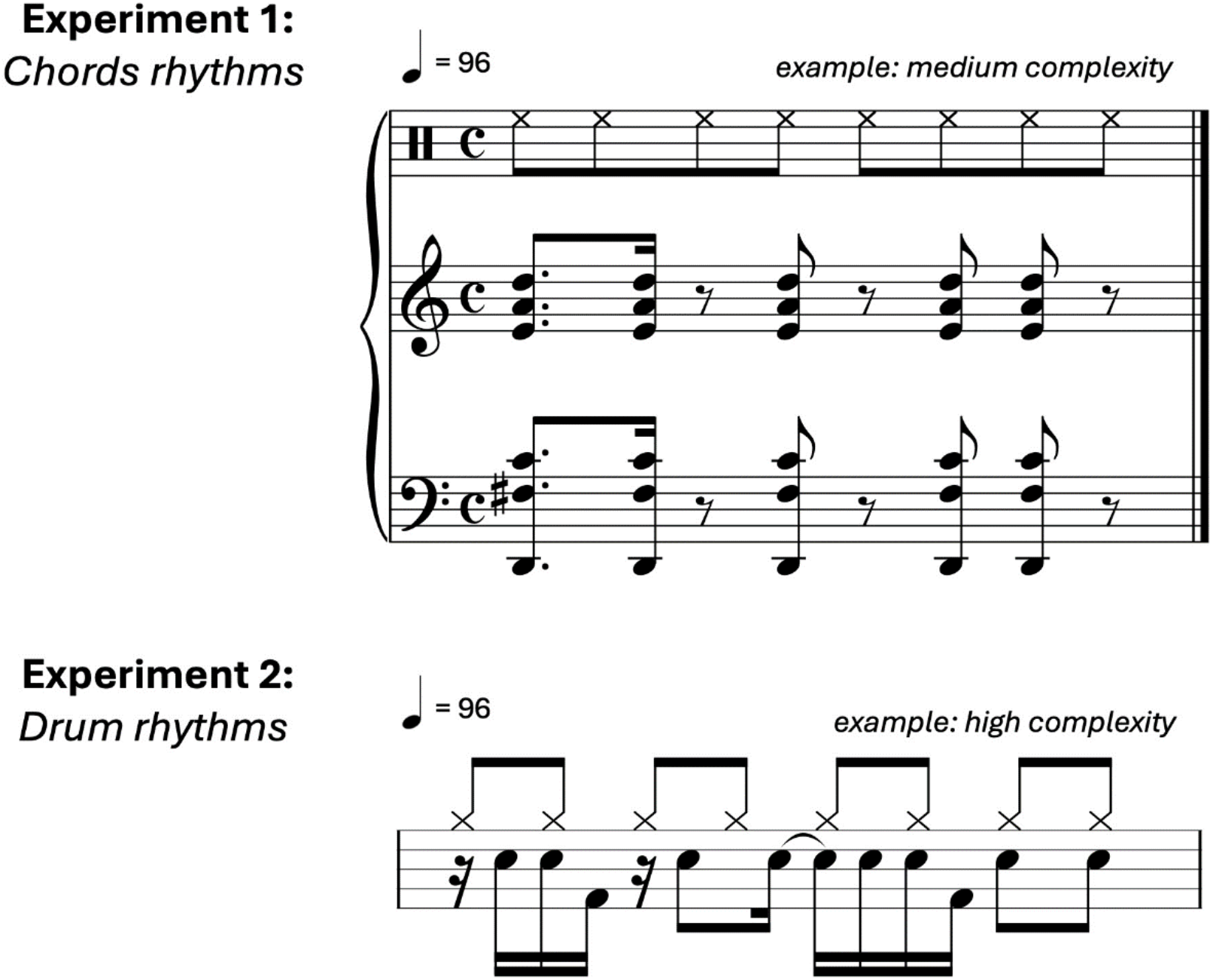
Example stimuli. *Top:* Chord-rhythm stimuli from Experiment 1, based on clave rhythms representing medium complexity in both harmony and rhythm. *Bottom:* Drum rhythm stimulus illustrating high rhythmic complexity. Both examples display a single bar.

**Figure 2.**
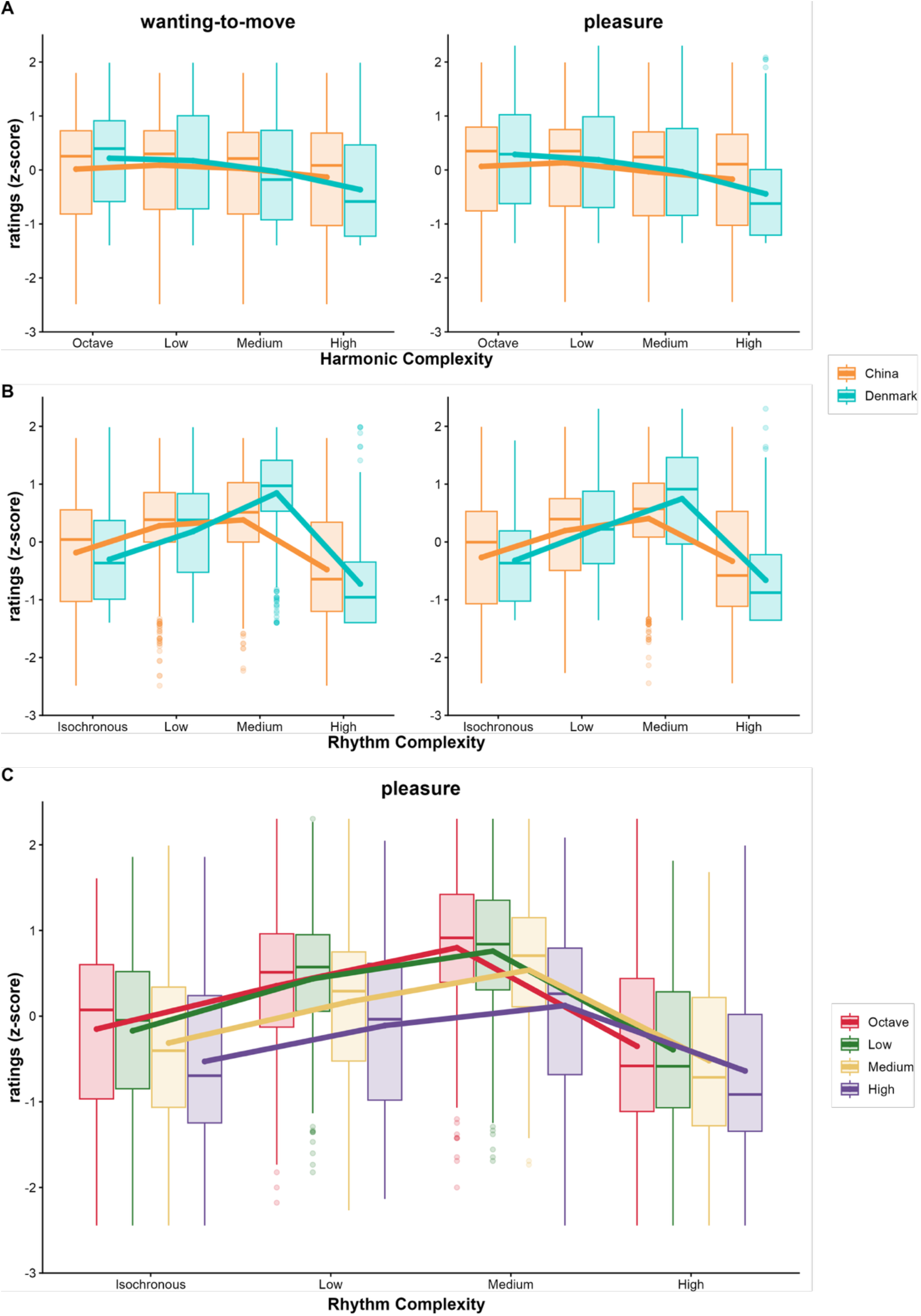
Interactions between harmonic complexity, rhythmic complexity, and nationality on wanting-to-move and pleasure ratings in response to chord rhythms. **(A)** Simple effects of the interaction between nationality and rhythmic complexity (left panel) or between nationality and harmonic complexity (right) on wanting-to-move ratings. (**B**) Simple effects of the interaction between nationality and rhythmic complexity (left panel) or between nationality and harmonic complexity (right) on pleasure ratings. (**C**) Simple effects of the interaction between rhythmic and harmonic complexity on pleasure ratings.

For the interaction between harmonic complexity and nationality, Chinese participants showed significantly smaller contrasts than Danish participants between the Medium and High levels (CN: *t*_(215)_ = 2.30, *p*_bonferroni_ = 0.067; DK: *t*_(215)_ = 7.35, *p*_bonferroni_ < 0.001; CN-DK: *t*_(215)_ = -3.59, *p*_bonferroni_ = 0.001). There was no significant group difference between the Octave and Low levels (CN: *t*_(215)_ = -1.46, *p*_bonferroni_ = 0.439; DK: *t*_(215)_ = 1.37, *p*_bonferroni_ = 0.521; CN-DK: *p*_bonferroni_ = 0.142), nor between the Low and Medium levels (CN: *t*_(215)_ = 0.92, *p*_bonferroni_ = 1.000; DK: *t*_(215)_ = 3.49, *p*_bonferroni_ = 0.002; CN-DK: *p*_bonferroni_ = 0.207). These results indicate that the relationship between harmonic complexity and wanting-to-move ratings does not follow an inverted-U shape. Instead, there appears to be a negative linear trend, where higher harmonic complexity is associated with reduced wanting-to-move ratings (**Figure 2A**, right panel). Additionally, ratings dropped more drastically at the higher end of complexity in Danish compared to Chinese participants.

#### Pleasure

Using *pleasure* ratings as the dependent variable (**Table S4**), we found significant interactions between nationality and rhythmic complexity (*F*_(3, 218.75)_ = 10.17, *q* < 0.001), nationality and harmonic complexity (*F*_(3, 239.19)_ = 9.53, *q* < 0.001), and rhythmic and harmonic complexity (*F*_(9, 2150.00)_ = 2.78, *q* = 0.019). The three-way interaction was not significant (*q* = 0.211).

Post-hoc analyses of group differences in contrasts between consecutive levels of rhythmic complexity revealed that Chinese participants had smaller contrasts than Danish participants between the Low and Medium levels (CN: *t*_(215)_ = -3.06, *p*_bonferroni_ = 0.008; DK: *t*_(215)_ = -7.65, *p*_bonferroni_ < 0.001; CN-DK: *t*_(215)_ = 3.27, *p*_bonferroni_ = 0.004), and between the Medium and High levels (CN: *t*_(215)_ = 8.00, *p*_bonferroni_ < 0.001; DK: *t*_(215)_ = 15.68, *p*_bonferroni_ < 0.001; CN-DK: *t*_(215)_ = -5.48, *p*_bonferroni_ < 0.001). There was no group difference between the Isochronous and Low levels (CN: *t*_(215)_ = -6.18, *p*_bonferroni_ < 0.001; DK: *t*_(215)_ = -8.19, *p*_bonferroni_ < 0.001; CN-DK: *p*_bonferroni_ = 0.447). These findings indicate an inverted-U relationship between rhythmic complexity and pleasure ratings, with medium rhythmic complexity evoking the highest pleasure and a more pronounced peak at the medium level for Danish compared to Chinese participants (**Figure 2B**, left panel).

Regarding harmonic complexity, compared to Danish participants, Chinese participants again showed smaller contrasts between the Medium and High level (CN: *t*_(215)_ = 2.33, *p*_bonferroni_ = 0.062; DK: *t*_(215)_ = 7.39, *p*_bonferroni_ < 0.001; CN-DK: *t*_(215)_ = -3.60, *p*_bonferroni_ = 0.001). There was no group difference between the Octave and Low levels (CN: *t*_(215)_ = -2.17, *p*_bonferroni_ = 0.092; DK: *t*_(215)_ = 1.17, *p*_bonferroni_ = 0.735; CN-DK: *p*_bonferroni_ = 0.058), nor between the Low and Medium levels (CN: *t*_(215)_ = 3.34, *p*_bonferroni_ = 0.003; DK: *t*_(215)_ = 3.85, *p*_bonferroni_ < 0.001; CN-DK: *p*_bonferroni_ = 1.000). Overall, the relationship between harmonic complexity and pleasure ratings did not follow the inverted-U pattern; instead, there was a negative linear trend for the pleasure ratings as harmonic complexity increased (**Figure 2B**, right panel). Similar to the urge-to-move ratings, ratings dropped more drastically at the higher end of complexity in Danish compared to Chinese participants.

For the interaction between rhythmic and harmonic complexity, specifically harmonic contrasts within rhythmic levels, significant effects were found only in Isochronous, Low and Medium levels of rhythmic complexity: Medium vs. High (Isochronous: *t*_(1422)_ = 3.52, *p*_bonferroni_ = 0.001; Low: *t*_(1422)_ = 3.97, *p*_bonferroni_ < 0.001; Medium: *t*_(1422)_ = 5.96, *p*_bonferroni_ < 0.001), and Low vs. Medium (Isochronous: *t*_(1448)_ = 2.96, *p*_bonferroni_ = 0.010; Low: *t*_(1448)_ = 2.82, *p*_bonferroni_ = 0.015; Medium: *t*_(1448)_ = 3.90, *p*_bonferroni_ < 0.001). All contrasts between the Octave and Low levels of harmonic complexity were not significant (*p*s_bonferroni_ = 1.000). At the High level of rhythmic complexity, all contrasts were not significant (*p*s_bonferroni_ > 0.133). Finally, there was a significant difference between the Medium and High levels of harmonic complexity in the contrast between the Medium and High levels of rhythmic complexity (Harmony_(Medium – High)_ × Rhythm_(Medium – High)_: *t*_(1935)_ = 2.92, *p*_bonferroni_ = 0.032). These results indicate that the effect of harmonic complexity on pleasure ratings was only present at lower and medium levels of rhythmic complexity, and disappeared at high rhythmic complexity. This suggests that rhythmic complexity may constrain the influence of harmony on affective responses (**Figure 2C**):

#### The role of musical sophistication

Additionally, there was a significant interaction between GMS and harmonic complexity (*F*_(3, 239.19)_ = 3.99, *q* = 0.044, **Table S4**), which was further tested through two approaches. First, a simple slope analysis revealed a significant positive trend of GMS on pleasure ratings for stimuli with medium harmonic complexity (*t*_(215)_ = 2.69, *p*_bonferroni_ = 0.008). There were no significant trends at the other levels (Octave: *p*_bonferroni_ = 0.060; Low: *p*_bonferroni_ = 0.914; High: *p*_bonferroni_ = 0.405) (**Figure S4A**).

We then conducted the pick-a-point analysis (**Figure S4B**), categorizing participants based on their GMS scores: those below the 10^th^ percentile were grouped as low, those between 45^th^ to 55^th^ percentile as medium, and those higher than 90^th^ percentile as high. Results showed that only in the low-GMS group, there were significant contrasts between the Octave and Medium levels (*t*_(55)_ = -3.07, *p*_bonferroni_ = 0.020), and between the Octave and High levels (*t*_(215)_ = -3.05, *p*_bonferroni_ = 0.021). No significant contrasts were observed between any two levels in the medium or high GMS groups (*p*s_bonferroni_ > 0.095).

### Experiment 2: Drum-Rhythms

#### Wanting to move

Using wanting-to-move ratings as the dependent variable, as shown in **Table S5**, the interaction between nationality and wSI was significant (*F*_(2, 364.91)_ = 13.88, *q* < 0.001). Model estimation showed that both the linear and quadratic terms of wSI significantly interacted with nationality (linear: β = 0.04, 95% *CI* = [0.02, 0.07]; quadratic: β = 0.08, 95% *CI* = [0.04, 0.11]).

Post-hoc analyses revealed that compared to Danish participants, Chinese participants exhibited a significantly smaller linear trend (CN: β = -0.10, 95% *CI* = [-0.16, -0.05]; DK: β = -0.19, 95% *CI* = [-0.24, -0.14]; CN-DK: *t*_(215)_ = 2.39, *p*_bonferroni_ = 0.018) and a significantly smaller quadratic trend (CN: β = -0.15, 95% *CI* = [-0.20, -0.10]; DK: β = -0.30, 95% *CI* = [- 0.35, -0.25]; CN-DK: *t*_(215)_ = 4.37, *p*_bonferroni_ < 0.001). These results indicate that the curvature of the inverted U-shaped relationship between wSI and wanting to move was flatter in Chinese than in Danish participants (**Figure 3A**). However, there was no differences in wSI at the peak of the curve between Chinese and Danish participants (95% *CI* = [-3.44, 4.28], **Figure 3B**).

**Figure 3.**
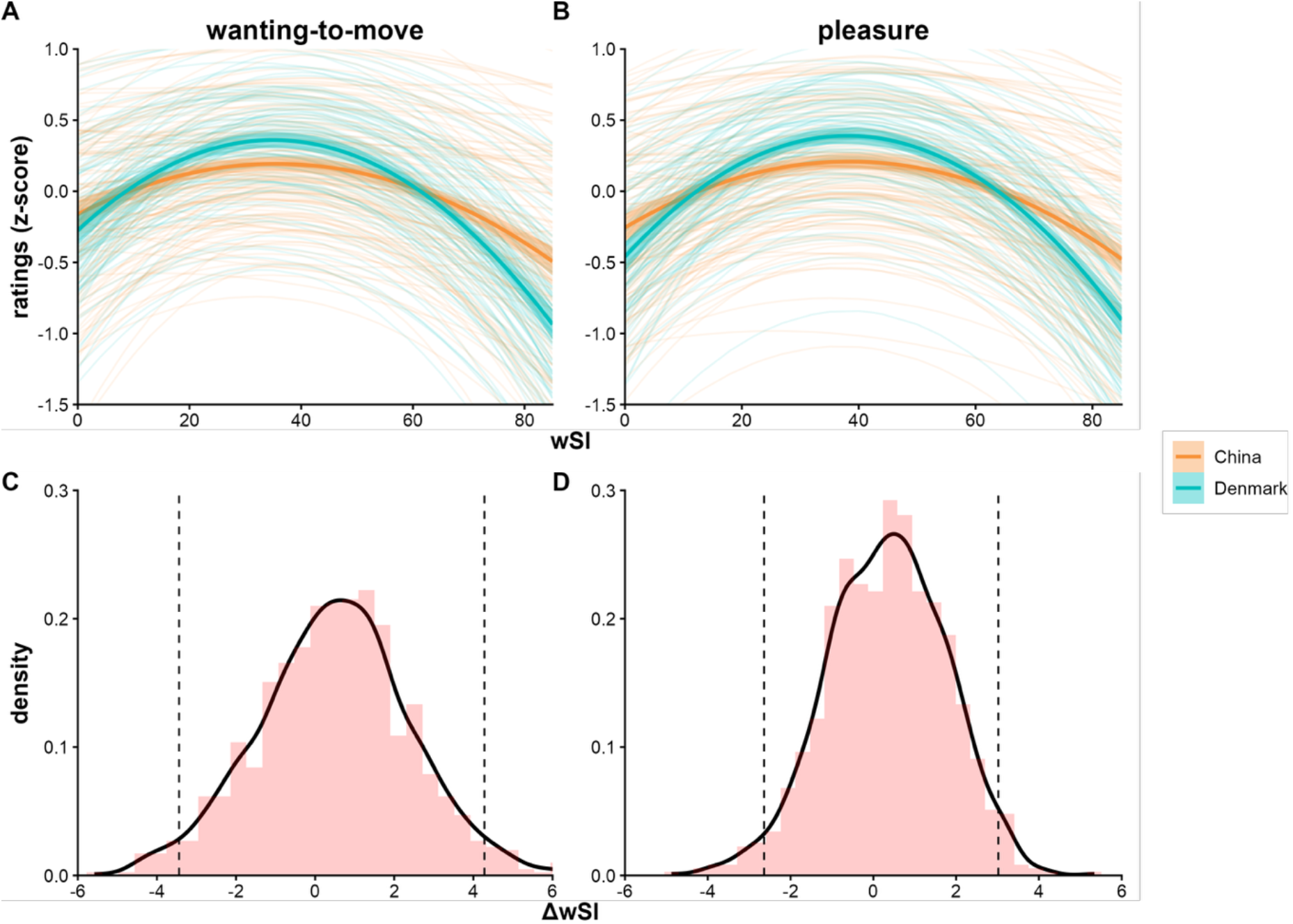
Interactions between wSI and nationality on wanting-to-move and pleasure ratings in response to drum rhythms. Interaction between wSI and nationality on wanting-to-move (**A**) and pleasure (**C**) ratings. Bootstrap results of wSI difference at the peak of the curve between Chinese and Danish participants for wanting-to-move (**B**) and pleasure (**D**) ratings.

#### Pleasure

Using pleasure ratings as the dependent variable, as shown in **Table S6**, a significant interaction between nationality and wSI also emerged (*F*_(2, 346.61)_ = 12.75, *q* < 0.001). Again, both the linear and quadratic terms of wSI significantly interacted with nationality (linear: β = 0.03, 95% *CI* = [0.00, 0.06]; quadratic: β = 0.08, 95% *CI* = [0.05, 0.12]).

Post-hoc analyses revealed that compared to Danish participants, Chinese participants had a significantly smaller quadratic trend (CN: β = -0.18, 95% *CI* = [-0.23, -0.13]; DK: β = -0.34, 95% *CI* = [-0.39, -0.29]; CN-DK: *t*_(215)_ = 4.63, *p*_bonferroni_ < 0.001). Although the linear trend was smaller in the Chinese group than in the Danish group, this difference did not reach significance (CN: β = -0.07, 95% *CI* = [-0.11, -0.02]; DK: β = -0.13, 95% *CI* = [-0.17, -0.08]; CN-DK: *t*_(215)_ = 1.74, *p*_bonferroni_ = 0.083). These findings suggest that the curvature of the inverted U-shaped relationship between wSI and pleasure was also flatter among Chinese than Danish participants (**Figure 3C**). However, there was no significant difference in the wSI value at the peak between the two groups (95% *CI* = [-2.64, 3.02], **Figure 3D**).

## Discussion

In this study, we investigated cross-cultural differences in affective responses to musical complexity, focusing on how rhythmic and harmonic complexities influence listeners’ ratings of pleasure and wanting to move among Chinese and Danish participants. Consistent with prior findings and our hypotheses, both cultural groups exhibited an inverted U-shaped relationship between rhythmic (but not harmony) complexity and affective responses. However, important cultural distinctions emerged in the curvature and strength of this relationship.

As for harmonic complexity, the absence of an inverted U-shape could reflect the fact that harmonies were presented within rhythmically varying contexts, where rhythm likely dominated affective responses.

### Cultural differences in rhythmic complexity responses

For rhythmic complexity, Danish participants showed a more pronounced inverted U response, with affective responses peaking at medium levels of complexity and dropping off at both lower and higher extremes. In contrast, Chinese participants showed a flattened response curve, with less differentiation across levels of complexity.

One plausible explanation for this cultural difference is variation in response styles. East Asian participants, including Chinese, are known to prefer mid-scale responses in self-report measures, whereas Western participants tend to use more extreme values ( Chen et al., 1995). However, this stylistic difference alone cannot fully account for the observed effects. A more substantive explanation involves culturally shaped differences in musical reward processing. As Wang et al. (2024) showed using BMRQ and network analysis, Chinese participants exhibit less cohesive clustering of sensory-motor musical reward features compared to North Americans, suggesting that music-induced movement may be a less salient or differently conceptualized form of musical pleasure in East Asian cultures.

The predictive coding framework offers a useful lens through which to interpret these findings. According to this model(Vuust et al., 2009, 2022) (Vuust et al, 2022), listeners rely on culturally acquired internal models to anticipate musical structure, and pleasure arises when those expectations are challenged in an optimal, moderately surprising way. Western enculturation may foster predictive models attuned to metrical regularity and deviations from this regularity (syncopations), thereby reinforcing pleasure and movement when rhythmic patterns strike a balance between predictability and surprise(Vuust et al., 2018) (Vuust et al., 2018). In contrast, Chinese musical tradition often emphasize expressive dynamics and fluid rhythmic structures rather than metrical rigidity (Wong, 2012; Mok, 1966). These traditions prioritize heterophony and the emergent texture of multiple melodic lines over the precise alignment of beats, leading to perceptual models that may downplay syncopation as a salient or rewarding musical feature. This broader cultural orientation may explain why Chinese participants responded less strongly to syncopated rhythms: such patterns may not optimally challenge their internal predictive models, resulting in a weaker emotional or embodied response.

Further supporting this view, prior research demonstrates that movement to music is culturally embedded, rather than a purely reactive phenomenon. For instance, South African participants in a cross-cultural study engaged significantly more bodily movements than Finnish participants, reflecting the integral role of dance in South African social and ritual activities (Himberg & Thompson, 2011). Similarly, it has been shown that movements to music are embedded in broader cultural concepts, not merely a spontaneous reaction to rhythm (Byczkowska-Owczarek, 2019; Pušnik, 2010). Applying this to our participants, Chinese students’ affective and embodied responses to rhythm may reflect a culture where public displays of spontaneous movement are more regulated or less common in everyday musical engagement.

The tightness-looseness cultural framework (Gelfand et al., 2011) may also offer an insight into the observed flattening of the inverted-U. China is often considered a relatively tight culture (Chua et al., 2019; Gelfand et al., 2011), emphasizing conformity and social regulation (Gelfand et al., 2020), which may dampen overt affective and physical responses to rhythmic complexity. In contrast, Denmark, often characterized as a looser culture, encourages more open and expressive forms of physical engagement with music, potentially amplifying pleasure responses at optimal complexity levels and contributing to the more distinct inverted U pattern.

### Cultural differences in harmonic complexity responses

In contrast to rhythmic complexity, harmonic complexity elicited relatively flatter inverted U-shaped responses in both cultural groups. While Danish participants exhibited steeper declines in ratings from low to high harmonic complexity, Chinese participants displayed greater affective responses especially to higher levels of harmonic complexity. This partially supports our hypothesis of relative heightened affective responses for the Chinese towards more complex harmonic structures.

(Matthews et al., 2019; Stupacher, Wrede, et al., 2022)(Witek et al., 2023)(Nettl, 2005)(Mok, 1966)These results are consistent with previous findings that speakers of tonal languages, such as Mandarin, exhibit heightened pitch perception abilities, evident in the higher prevalence of absolute pitch (Deutsch et al., 2004; Pfordresher & Brown, 2009) and superior performance in melodic tasks (Arunkumar & Sadakata, 2021; Joe et al, *in prep)*. Chinese listeners, even those without formal music training, often demonstrate heightened sensitivity to fine-grained pitch differences (Liu et al., 2021; Ngo et al., 2016), a capacity that supports a deeper appreciation of complex harmonic structures. Similarly, research has shown that both tonal language speakers and musicians possess enhanced auditory encoding of musical pitch (Bidelman et al., 2013), and musicians show peak affective responses at higher levels of harmonic complexity than non-musicians (Witek et al., 2023). Our results align with the idea that lifespan enculturation—whether musical or linguistic—can shape affective responses to harmonic complexity through implicit learning of pitch structures.

### Interaction of harmony and rhythm in affective responses to music

In line with our hypothesis, we observed cultural differences in how rhythmic and harmonic complexity interact to shape affective responses. However, these differences did not manifest clear shits in peaks of inverted-u relationships. Rather, responses specifically to harmonic complexity were, for the Chinese, characterized as flatter without an explicit peak, while the Danish participants showed a more negative sloped relationship across the different levels of harmonic complexity. In contrast, rhythmic complexity elicited a clearer inverted U-shaped pattern in both groups, though with a flatter curvature for the Chinese participants. One possible explanation to why complexity of rhythm and harmony elicit different response patterns, is that harmony and rhythm interact to shape affective responses to music in culturally dependent ways. Matthews et al. (2019) demonstrated that harmonic properties influence wanting to move indirectly through its impact on pleasure, whereas rhythm directly affects both wanting to move and pleasure. Given Chinese listeners’ enculturated higher sensitivity to pitch, it is plausible that harmonic context amplifies their pleasure responses to rhythm, thereby elevating their experience of wanting to move.

### Universal mechanisms in affective responses to music

Despite cultural distinctions in curvature, it is noteworthy that both Chinese and Danish participants displayed inverted U-shaped responses to rhythmic complexity. This convergence supports the existence of universal, low-level mechanisms underlying affective engagement with rhythm. For example, moderate levels of syncopation may optimally engage predictive processing and learning mechanisms to maximize motor engagement and reward (Friston, 2010; Margulis, 2014; Matthews et al., 2023). While cultural background shapes internal models and aesthetic preferences, the core cognitive mechanisms of predictive processing appear transcending cultural boundaries (Millidge et al., 2022).

### Limitations and future perspectives

Despite the novel insights provided by this study, several limitations should be considered. First, the sample was restricted to Chinese mandarin speakers and Danish participants, limiting generalizability. Future research could benefit from including a broader range of cultures to capture a fuller spectrum of cultural variability. Second, the stimuli could be enhanced by incorporating more diverse harmonic progressions and cadences, allowing for a finer-grained examination of how harmonic evolution interacts with rhythm and culture in shaping affective responses. In particular, examining harmonic complexity in isolation, without concurrent rhythmic variation, would help clarify how cultural differences specifically influence affective responses to harmony. Third, cultural familiarity with the rhythmic patterns used in the study remains a potential confound. Chinese participants may have found the syncopated rhythms less familiar, leading to muted responses. Future studies could systematically manipulate familiarity or include culturally indigenous rhythms to better isolate effects of complexity from those of exposure and preference. Fourth, individual differences in musical sophistication were not fully independent of culture, as Chinese participants scored higher on average GMS than Danish participants. Consequently, the grouping of participants into high, medium, and low GMS categories may have partially reflected cultural background, potentially confounding effects of musical sophistication with those of cultural exposure.

## Conclusion

In conclusion, this study demonstrates that both universal and culture-specific mechanisms shape how musical complexity influences affective responses. While both Chinese and Danish participants exhibited inverted U-shaped responses to complexity, significant cultural differences emerged in the sensitivity and curvature of these relationships. Danish participants showed heightened sensitivity to rhythmic complexity, whereas Chinese participants displayed flatter responses to rhythmic complexity along with heightened appreciation for complex chords, likely reflecting linguistic pitch experience and cultural musical exposure. These results suggest that while universal mechanisms in music appreciation exist, cultural influences significantly modulate how musical complexity is appraised. Our findings advance our understanding of how culture shapes musical perception and underscore the need for culturally inclusive models of musical cognition.

