## Supplementary Information for "Universality Meets Cultural Nuances: Affective Responses to Rhythmic and Harmonic Complexity Across Cultures"

***Response style bias***

**Commented [YD1]:** Move all methods and results regarding response style bias to SI, just leaving one summary sentence in the main manuscript.

To assess potential cross-cultural response style bias, we conducted complementary distributional analyses in both experiments. In Experiment 1, we examined full rating distributions across all levels of harmony (figure S5) and rhythm (figure S6) complexity, whereas in Experiment 2, we performed a representative-stimulus analysis using subsets of stimuli drawn from the lowest, middle, and highest levels of wSI (four stimuli per level) (figure S7).

In Experiment 1, across both pleasure and wanting-to-move ratings, participants in both cultural groups generally exhibited non-unimodal (predominantly bimodal) response distributions across complexity levels. However, this pattern was not fully symmetric across groups. While Chinese participants showed relatively stable bimodal distributions with balanced peaks across conditions, Danish participants displayed a systematic deviation at high complexity in both harmony and rhythm. Specifically, their distributions became strongly left-skewed, with a pronounced accumulation of responses near the lower bound of the scale and a reduced or attenuated high-rating mode, rather than a symmetric bimodal pattern with a clear central saddle point. At lower and intermediate complexity levels, the two groups showed more comparable bimodal structures, and the saddle point between modes, where present, aligned closely with the z-scored midpoint of the rating scale.

In Experiment 2, the representative-stimulus analysis yielded a largely consistent picture across groups. Both Danish and Chinese participants again showed clear bimodal distributions across pleasure and wanting-to-move ratings. The distributions at the medium complexity level were highly similar between groups, including a relatively stronger high-rating mode. Group differences were primarily evident at the high complexity level: Chinese participants maintained relatively consistent distributional shapes across all levels, whereas Danish participants again showed a shift toward lower ratings. However, unlike in Experiment 1, this shift did not manifest as a pronounced boundary accumulation, but rather as a relative increase in the low-rating mode compared to the high-rating mode.

Across both experiments, these distributional characteristics do not match patterns typically associated with canonical response style biases, such as uniform scale compression or expansion (e.g., global restriction or indiscriminate use of extreme response categories). Instead, the observed differences are condition-specific—most notably at high complexity—and qualitatively distinct between groups. Taken together, these findings suggest that the reported effects are unlikely to be driven by cross-cultural response style bias, but instead reflect genuine differences in how high-complexity stimuli are evaluated across cultural groups.

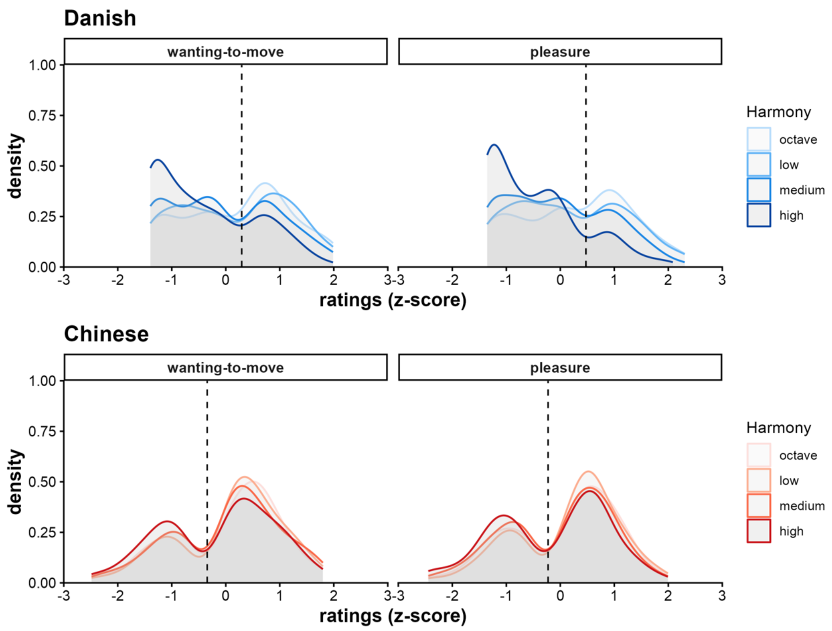


Figure S5. Distribution of wanting-to-move and pleasure ratings under different levels of harmony complexity in Experiment 1. Black dashed vertical lines indicate the z-score corresponding to the midpoint of the rating scale (50/100).

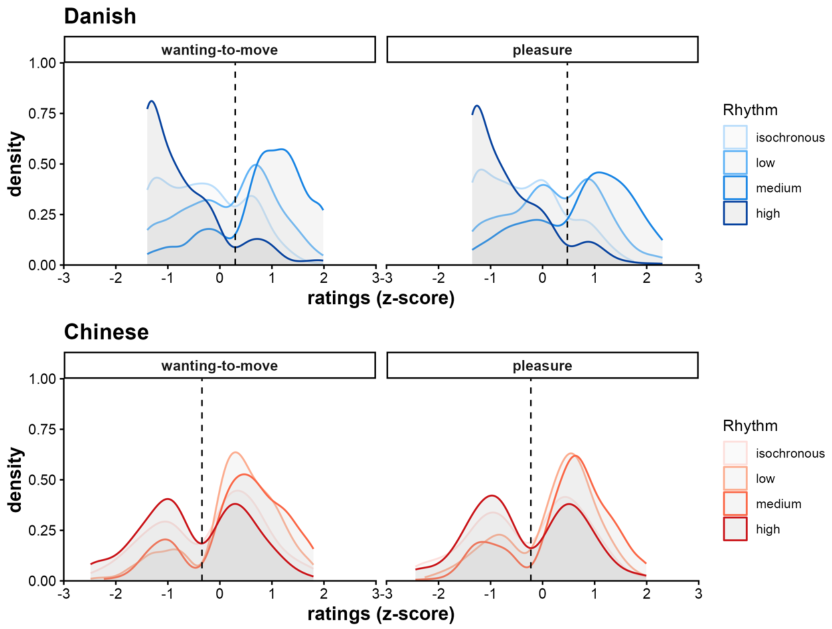

Figure S6. Distribution of wanting-to-move and pleasure ratings under different levels of rhythm complexity in Experiment 1. Black dashed vertical lines indicate the z-score corresponding to the midpoint of the rating scale (50/100).

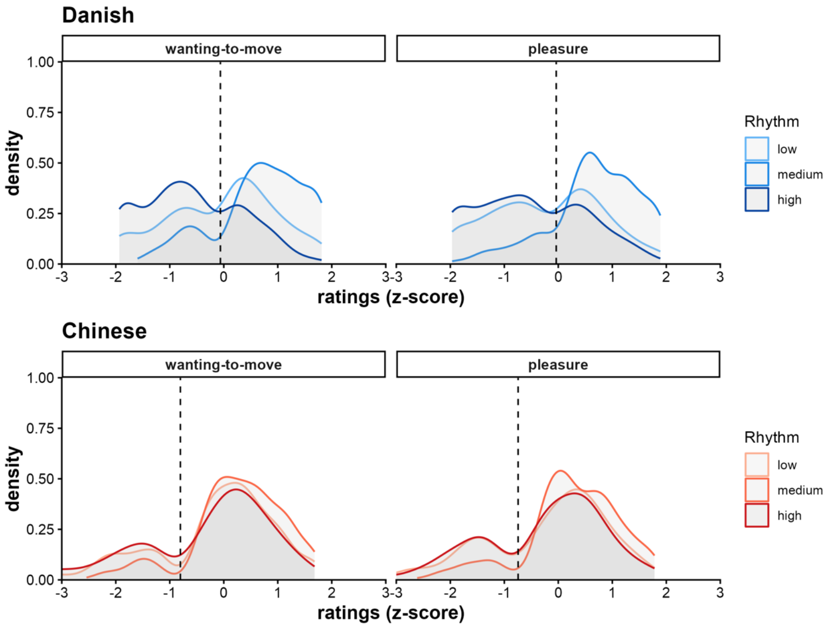


Figure S7. Distribution of wanting-to-move and pleasure ratings under different levels of rhythm complexity in Experiment 2. Each distribution curve represents ratings aggregated from a subset of four representative stimuli (out of 24) for each complexity level. Black dashed vertical lines indicate the z-score corresponding to the midpoint of the rating scale (50/100).
